# Size in the city: morphological differences between city and forest great tits have a genetic basis

**DOI:** 10.1101/2025.02.05.636678

**Authors:** Barbara M. Tomotani, Mika Couweleers, Bram ten Brinke, Anne Walboom, Kees van Oers, Marcel E. Visser

## Abstract

Animals living in cities are smaller than their conspecifics from rural areas but whether such differences are caused by genetic differences or food constraints remains untested. We performed a multi-generation common garden study where we raised great tits (*Parus major*), originating from eggs collected from multiple Dutch cities and forests under the same conditions for two generations. Offspring from city birds had a smaller tarsus than forest birds in both generations, demonstrating that these morphological differences are genetic. Next, we tested whether size differences are an adaptation to the low food abundance when offspring are raised in the city. Third generation birds of both origins were given food amounts mimicking being raised in forests or cities during the second part of their nestling development. While the treatment resulted in birds in the lower feeding frequency treatment to be smaller, city and forest birds responded the same way, suggesting that city birds do not cope better with reduced food availability. Our study shows that the smaller size of urban birds has a genetic basis and is not only caused by a plastic response to restricted resources in the urban environment. Our experiment does not provide evidence that these genetic differences have evolved as an adaptive response to a reduced food availability in cities.

## Introduction

Cities are rapidly expanding across the globe, with estimations that two thirds of civilians will live in urban areas by 2050 (United Nations, 2018). Urbanisation is considered a major threat for wildlife (Hendry *et al*., 2017) but may also play a role as an evolutionary driver of adaptation (Johnson *et al*., 2018). Cities are known for their strongly modified environments exposing organisms to a number of distinct challenges when compared to natural habitats (Isaksson, 2018; Dominoni *et al*., 2020; Sanders *et al*., 2021; Thompson *et al*., 2022; Tomotani *et al*., 2023). Morphological differences, in particular, are consistently reported between city and forest animals. In birds, many studies have shown a lower body mass and/or smaller skeletal size of city birds (Liker *et al*., 2008; Meillère *et al*., 2015, 2017; Biard *et al*., 2017; Caizergues *et al*., 2018a, 2021; Senar & Björklund, 2021; Thompson *et al*., 2022; Santos *et al*., 2023), but differences vary in magnitude depending on the geographical location (Evans *et al*., 2009) and species (Brouwer *et al*., 2024).

The smaller size of city birds would normally be attributed to the harsher city life, where food is less abundant, particularly when the bird is still developing (Seress *et al*., 2020). Since tarsus length of songbirds becomes fixed at an early time during the development, poor conditions experienced early in life may carry-over into adulthood impacting survival and breeding (Lindström, 1999; Cleasby *et al*., 2011; Caizergues *et al*., 2021). Despite an apparent abundance of food in cities, especially due to the popularization of garden feeding (Crates *et al*., 2016), the availability of high quality food for the offspring still seem to be limited (Seress *et al*., 2020), as the food requirements of offsprings may differ from those of the adults. For example while great tit chicks depend on caterpillars as their a main food source (Naef-Daenzer *et al*., 2000) adult great tits have a mixed insect and seeds diet in winter (Wansink & Tinbergen, 1994) and can benefit from the food provided in gardens. Indeed, birds in the city are reported to feed their offspring with different food items when compared to forests, often of poorer quality or in smaller quantities than forest birds (Seress *et al*., 2012; Sinkovics *et al*., 2021). For example, the amount of caterpillars in the diet of great tit chicks is reported to be lower in cities than forests (Seress *et al*., 2018; Sinkovics *et al*., 2021). Furthermore, experimental studies where food supplementation is offered to both city and forest great tits have shown that food may indeed be a limiting factor in the breeding success of city birds, because food supplemented city chicks grow almost as much as forest chicks (Seress *et al*., 2020).

Recent investigations, however, have shown that the phenotypic divergence between urban and rural populations may also have a genetic basis (Reid *et al*., 2016; Campbell-Staton *et al*., 2020). For example, heat tolerance is genetically higher in urban than forest anole lizards (Campbell-Staton *et al*., 2020). This is in line with the finding that evolutionary adaptations in natural populations can occur within short timescales, particularly in response to human activities (Hendry *et al*., 2008; Bonnet *et al*., 2022). When adult individuals living in distinct environments are compared, it is not possible to unequivocally separate genetic and environmental effects, such as those experienced early in life. Thus, to identify the origin of the size differences between city and forest birds, we need an experimental approach.

Here we used a classical “common garden” experiment set-up, where we collected eggs from multiple city and forest populations. These eggs hatched, and the offspring that arose from them were hand raised in a common location (same early-life environment) as well as phenotyped under the same condition (same adult environment) (de Villemereuil *et al*., 2016). To minimize maternal environmental effects, we repeated this for two generations under the same conditions. Then, after rearing birds from city and forest populations that differ in body size (Fig. 1) under same conditions for two generations, we tested whether a smaller size in city birds could be an advantage when being raised under conditions of low food availability. We predicted that city birds would grow better compared to forest birds under lower food availability, while forest birds would grow better when food is more abundant. For this we manipulated the early life feeding conditions and measured the effects on body size for birds originating from urban and forest populations.

**Figure 1:**
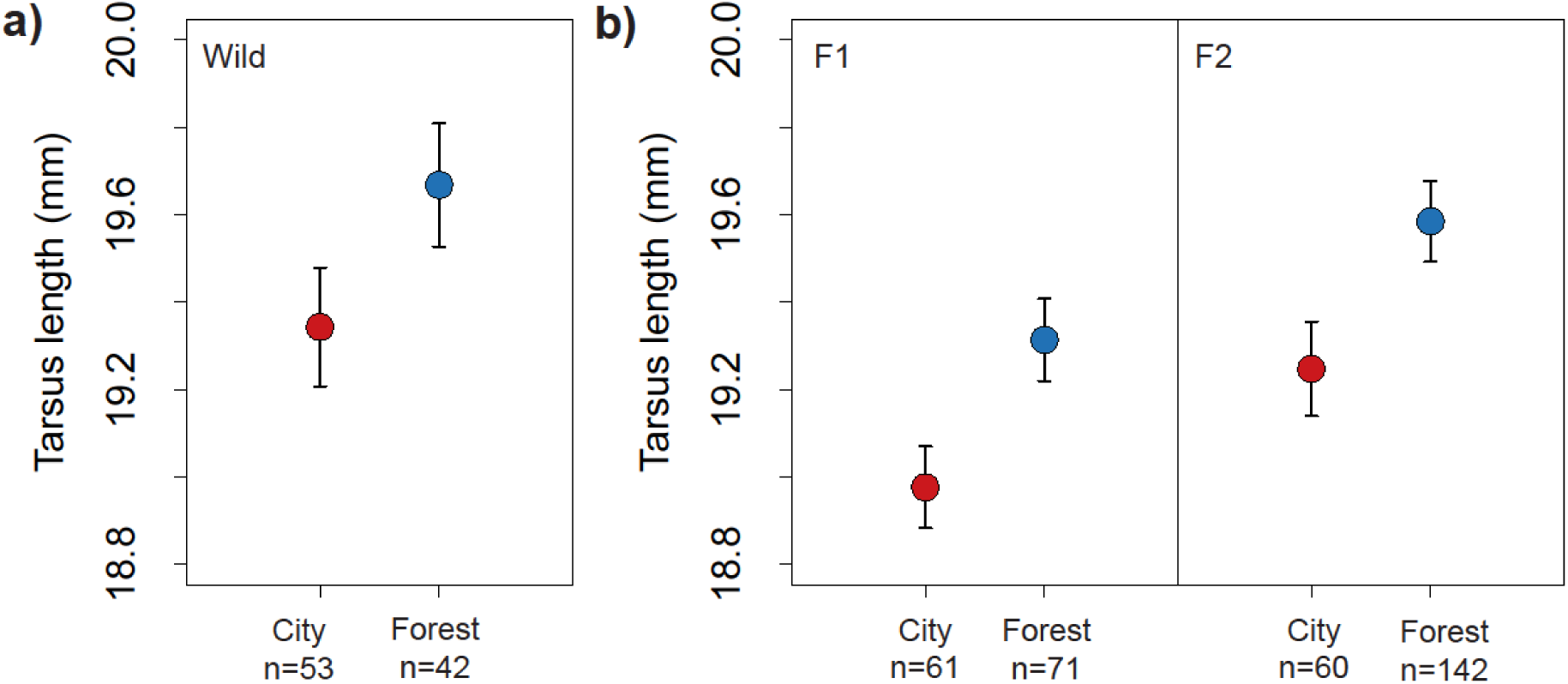
Differences in tarsus length between city and forest birds: **a)** When comparing wild caught city (Utrecht) and forest (Heikamp) birds. **b)** When comparing birds in common garden (F1 and F2 generations). Symbols: marginal means from the models containing the additive effects of **a)** site, sex and year and **b)** origin, sex, generation, centered hatching date^2 and centered hatching date. Bars: standard errors.

## Methods

See the supplementary Table S1 for the detailed sample sizes for each part of this study. All experimental procedures in the lab and in the field were carried out under licenses of the Central Authority for Scientific Procedures on Animals (Project AVD 80100 2019 9005) and the Animal Welfare Body (IvD) of the Royal Netherlands Academy of Sciences (KNAW; protocols NIOO21.04 and NIOO23.07.

### Experimental procedure

#### Wild birds comparison

During the spring of 2020, 72 breeding birds were caught in their nest boxes (as part of another experiment (Tomotani *et al*., 2023)) in the city of Utrecht (lat: 52.090, long: 5.121) and the forest of Heikamp (lat: 52.031, long: 5.834), just next to the Hoge Veluwe National Park (lat: 52.041, long: 5.856) where eggs for the common garden experiment also came from (see below). The birds had their tarsus measured (nearest 0.1 mm) and were further recaptured and remeasured on other occasions, resulting in 95 measurements that we used here for a city/forest comparison of tarsus length.

#### Common garden experiment

##### First generation (F1)

In spring 2021, we carried out a common garden experiment where birds from distinct populations were raised from eggs under the same environment. We used great tit populations in four Dutch cities (Amsterdam lat: 53.372, long: 4.893, Delft lat: 52.012, long: 4.359, Rotterdam lat: 51.92, long: 4.48 and Utrecht) and five Dutch forests (Bennekomse Bos, lat: 52.003; long: 5.708, Hoge Veluwe, Liesbos lat: 51.582, long: 4.700, Oosterhout lat: 51.873, long: 5.584 and Vlieland lat: 53.295, long: 5.051).

During spring 2021, we selected 5-8 first broods from each population with non-extreme first egg laying dates. The difference in first egg dates was on average five days from the population mean laying date. We then collected five eggs per brood and immediately transferred them to a foster brood in the forest of Bennekomse Bos. Chicks hatched in the foster nest and were raised by their foster parents until 10 days after hatching, when they were taken to the

Netherlands Institute of Ecology (NIOO-KNAW) bird facilities to be hand raised. This resulted in 136 chicks (65 city, 71 forest). In all analyses we only included birds that made it to the independence stage and were thus fully grown and their tarsus could be measured, thus here 131 (61 city, 71 forest). More details about egg collection and transport can be found in (Visser *et al*., 2025), all eggs in this study were transport by car.

Hand raising was done according to standardized protocols (Drent *et al*., 2003). In brief, upon arrival at NIOO-KNAW, nestlings were transferred to a wooden box in groups of three to four birds, each box containing three compartments with a natural parasite-free nest in each. When birds fledged, at around 17–20 days after hatching, they were transferred to small a wire mesh cage in groups of three. Birds are independent around day 35 after hatching, after which they were moved to standard individual cages of 0.9 m× 0.4 m× 0.5 m. Birds were always kept under natural light conditions in acoustic and visual contact with each other.

##### Second generation (F2)

In early January 2022 we formed breeding pairs using the F1 birds, pairing individuals from the same populations (14 city pairs and 25 forest pairs). These pairs were kept in aviaries (4 m x 2 m x 2 m or 2 m × 2 m × 2.25 m, l x w x h) and allowed to breed in spring. Due to the low number of males and females obtained, we did not make breeding pairs of the Bennekomse Bos and Liesbos populations. Eggs were collected directly after laying and moved to a foster brood in the forest of Bennekomse Bos. Chicks hatched in the foster parent nest and were raised by their foster parent until 10 days after hatching, when they were taken to the NIOO-KNAW facilities to be hand raised according to the standardized conditions described above. We obtained 207 chicks (63 city, 144 forest), of which 202 (60 city, 142 forest) reached independence and could be included in the analyses.

#### Feeding experiment

In early January 2023 we formed breeding pairs using the F2 birds, pairing individuals from the same populations (10 city and 20 forest pairs), those were kept in aviaries (4 m x 2 m x 2 m, l x w x h) and allowed to breed in spring. Due to the availability of breeding pairs, we only used birds from the Hoge Veluwe as our forest population. Eggs were collected in the breeding aviaries and again moved to a foster brood in the forest of Bennekom. Chicks from this third generation (F3) were raised by their foster parent until 10 days old when they were taken to the NIOO-KNAW to be hand raised. This time, however, chicks were fed following an experimental feeding regime so, when they arrived at the institute, they were divided into three treatments: food every 30, 45 and 60 minutes (with 30 minutes being the standard feeding rate as also used for the F1 and F2 birds) from the moment they arrived (10 days old) until fledging. We obtained 106 chicks (34 city, 72 forest) that were divided between the treatments, of which 100 (34 city, 66 forest) could be included in the analyses (see Supplementary Table S1).

#### Phenotyping

All chicks (F1, F2 & F3) had their tarsus measured (nearest 0.1 mm) by the same observer when fully grown.

#### Egg mass

In addition to the tarsus measurements, we also weighed a total of 1864 eggs (precision 1 mg), laid by the females in the wild (by the wild generation P: 77 city and 109 forest) and in common garden (by the first generation F1: 509 city and 619 forest; by the second generation F2: 123 city and 425 forest).

### Data analyses

Analyses were carried out in R (4.3.0) (R core team, 2023). Unless stated otherwise, we used linear mixed effect models (package “lme”, “lmer” function). P-values were obtained via backwards selection by dropping the terms of interest in each step and comparing the models via a Kenward Roger approach for approximation of F-values (package “pbkrtest”, “KRmodcomp” function).

#### Wild birds comparison

We tested whether tarsus length differed between city and forest birds. We included site (city or forest), sex (M/F) and the interaction between them as explanatory variables. Because some birds were measured multiple times throughout their lifetime, we also included year as fixed effect and individual as random effect.

#### Common garden experiment

We compared whether tarsus length differed between birds with a genetic city or forest origin using birds from the F1 and F2 generations. We included origin (city or forest), generation (F1/F2) and sex (M/F) as fixed effects and also included hatching date and hatching date squared (henceforth: HD2) as fixed effects in order to correct for differences in the foster brood condition. Because the peak and spread of hatching dates differed significantly between the two years when the experiment took place (see Supplements 1 and 4) values were centered around the mean hatching date per year. To take into account potential genetic differences of family or population of origin, we included family nested within population as random effects. We also included the interactions between origin and HD2, hatching date, sex and generation. In the model selection procedure, we dropped the interactions and main effect of origin for obtaining the p-values, while retaining the terms sex, generation and hatching date/HD2 in the model as nuisance variables.

#### Feeding experiment

With the experiment, we were interested in testing whether birds from forest- or city origin would respond differently to the feeding treatment in terms of their size, thus whether there was an interaction between origin and treatment. We fitted treatment (30 min, 45 min or 60 min), origin (city or forest) and sex (M/F) as fixed effects and also included the hatching date and HD2 as fixed effects as in the previous analysis. Once more, to account for potential genetic differences of family or population of origin, we included family nested within population as random effects. Apart from the interaction between origin and treatment we also included the interactions between origin and hatch date, origin and sex, treatment and hatch date and treatment and sex. In the model selection procedure, we dropped the interactions and main effect of origin and treatment for obtaining the p-values, while retaining the terms sex, generation and hatching date/HD2 in the model as nuisance variables.

#### Egg mass

We compared if the egg mass differed between city and forest birds across generations. We included site (city or forest), generation (P, F1 or F2) and the interaction between them as explanatory variables. To take into account potential genetic differences of family or population of origin, we included the mother identity nested within population and the father identity nested within population as random effects.

## Results

### Wild birds comparison

Similarly to what is reported in other populations (Caizergues *et al*., 2021), we found a significant difference between the tarsus length of city and forest birds (*F1,48.18* = 5.76, *p* = 0.01 Fig. 1a), with city birds being smaller than forest birds. In addition, males had longer tarsi than females (*F1,41.17* = 20.27, *p* = 0.01).

### Common garden experiment (F1 and F2 birds)

Birds originating from city families had a significantly smaller tarsus than birds from forest families in both generations (*F1,5.67* = 9.02, *p* = 0.03, city estimate = 18.86 ±0.10mm, forest estimate = 19.20 ±0.09mm, Fig. 1b).

### Feeding experiment (F3 birds)

We tested whether birds from a forest origin grow less well under a restricted food regime compared to city birds, which would suggest that being small is an advantage when exposed to low food availability in cities. Thus, we tested if birds from different origins (city or forest families, offspring from the F2 birds above) differed in their size when fed every 30, 45 or 60 minutes between day 10 post-hatching and fledging. There was a significant effect of treatment on tarsus length (*F2,80.01* = 3.91, *p* = 0.02, Fig. 2), with birds in the 60 min treatment being smaller than birds in the 45 and 30 min treatments (estimates 30 min = 19.54 ±0.25mm, 45 min = 19.54 ±0.25mm, 60 min = 19.25 ±0.24mm). However, birds originating from city or forest families did not differ in their sensitivity to the food treatments (no significant interaction between origin and treatment; *F2,74.12* = 0.07, *p* =0.93), and thus city and forest birds developed similar tarsus differences between the treatments (Fig. 2). There was also no main effect of origin, thus no overall difference in tarsus length between city and forest birds (F1, 1.19 = 0.00, p = 0.98).

**Figure 2:**
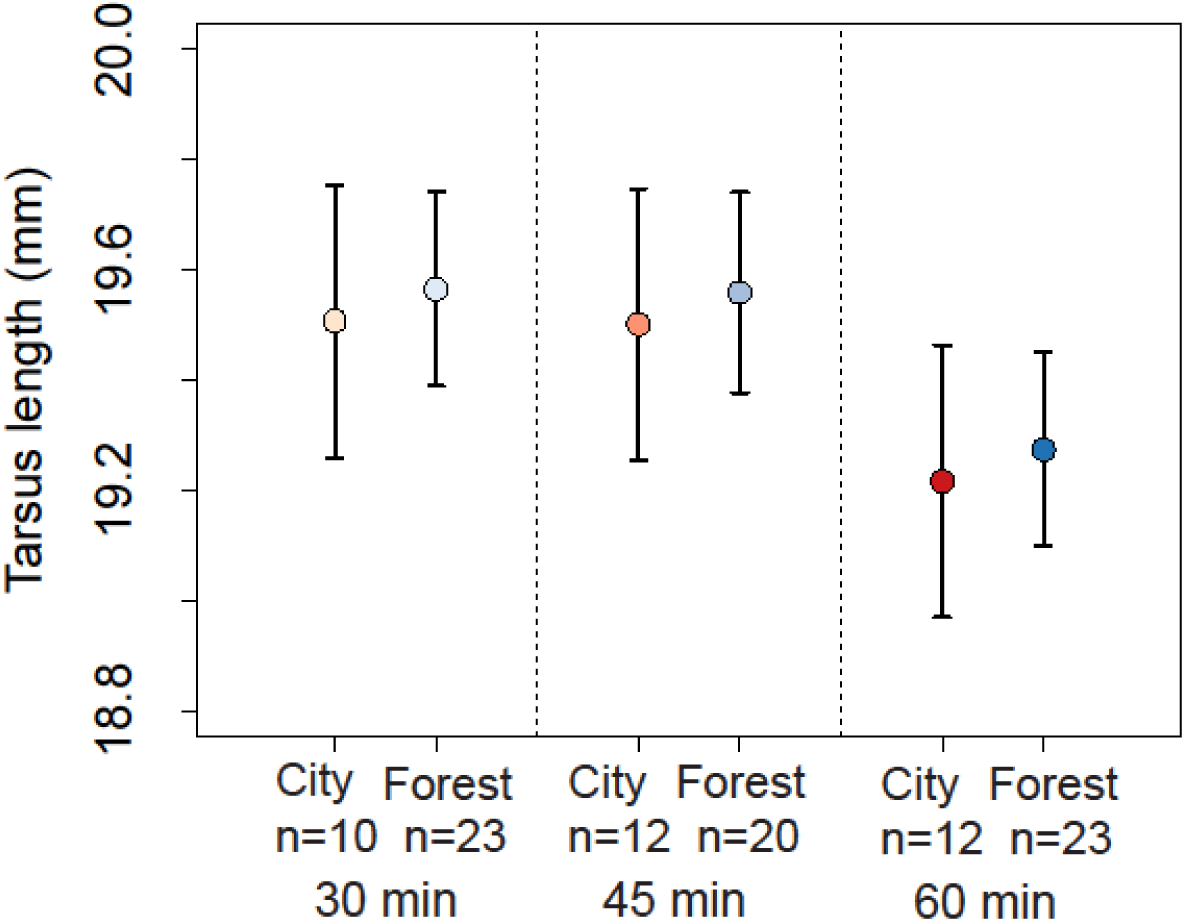
Differences in tarsus length of F3 birds. Symbols: marginal means from the models containing the additive effects of treatment, origin, sex, generation, centered hatching date^2, centered hatching date and the interaction between hatching date and origin. Bars: standard errors.

### Egg mass

There was a non-significant interaction between mother origin and mother generation (*F2,80.77* = 2.94, *p* = 0.06). A post-hoc analysis showed a significant difference between city and forest eggs laid in the wild by the P generation (t = -3.14, *p* < 0.01) but not between eggs laid in common garden by the F1 and F2 generation (F1: t = -1.05, *p* = 0.31; F2: t = 0.39, *p* = 0.70; Fig. 3).

**Figure 3:**
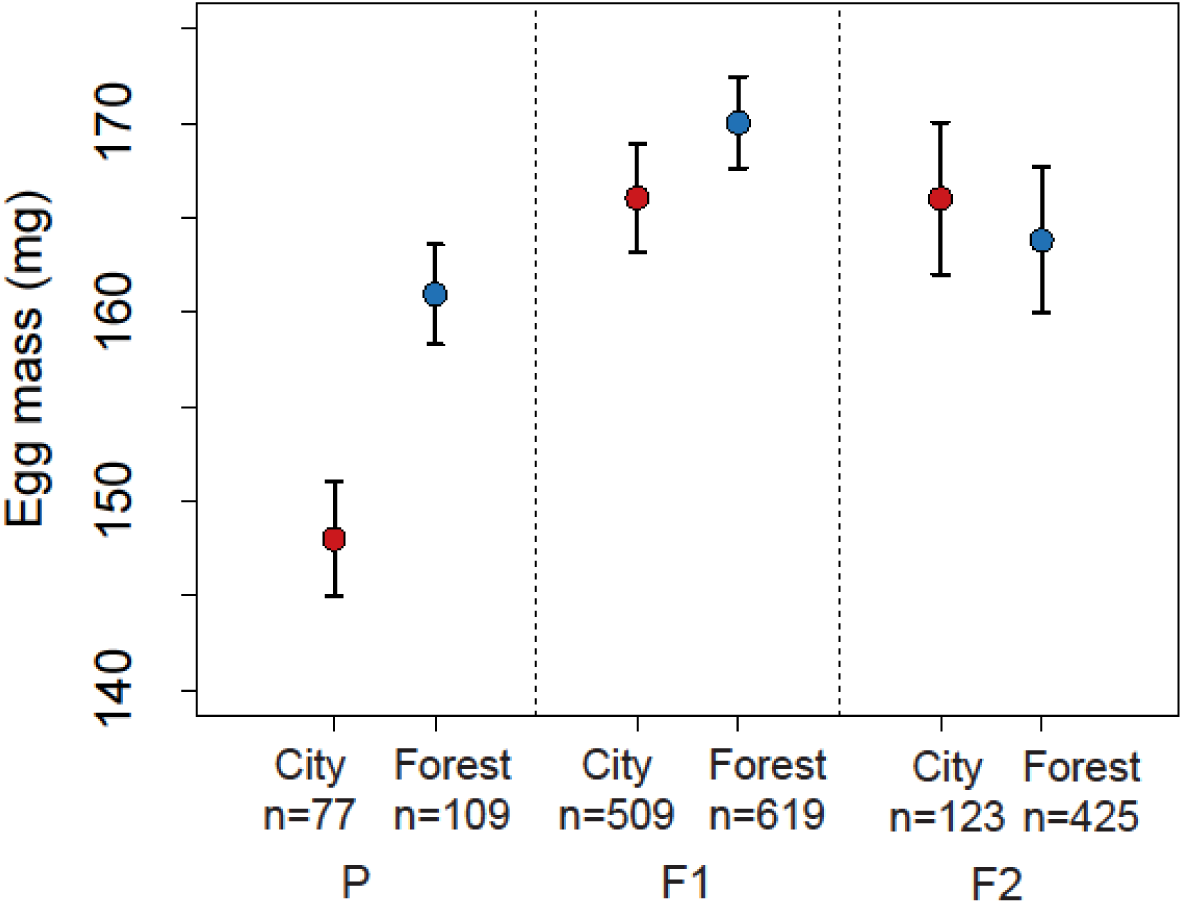
Differences in egg masses laid by mothers from P, F1 and F2 generations from city or forest origins. P eggs were laid in the wild, F1 and F2 in common garden. Closed symbols: marginal means from the models containing the effects of mother site, mother generation, the interaction between mother site and mother generation and centered laying date. Bars: standard errors; open symbols: raw measurements.

## Discussion

Food is regularly suggested to be the main cause of size differences between urban and forest birds where the smaller body size would be caused by the reduced food quality or amount found in cities (Seress *et al*., 2020; Caizergues *et al*., 2021). Indeed, experiments have shown that supplementing nestlings with food is partly able to compensate for differences between city and forest birds (Seress *et al*., 2020). But in this study, we experimentally show that differences in body size found in the wild between city and forest birds persist even after two generations. This shows that city and forest great tit populations differ in their genetic makeup: urban great tits are genetically smaller. To our knowledge, this finding had not been demonstrated experimentally so far. In a very recent study on a (single generation) common garden experiment with French great tits from an urban-rural gradient birds originating from an urban population were also shown to be smaller albeit, more due to a lighter body mass than shorter tarsus (Thompson *et al*., 2025). Nevertheless, this indicates that the genetic basis of a smaller size in city birds is not limited to Dutch birds.

Then, we explored the possibility that the body size of city birds is less compromised by lower food availability than city birds. Although the differences in tarsus length seem small, offspring body size is known to have an impact on the survival probability of chicks (Rodríguez *et al*., 2016), and even have indirect effects on the probability of fledgling via competition among siblings (Oddie, 2000). If a smaller body size allows birds to better cope with food availability in the city, this would result in a different city/forest nestling growth under distinct feeding regimes. The results of our experiment, however, did not support the hypothesis that a smaller body size would be beneficial when food is scarce.

Measuring birds raised in common conditions over two generations and not just raising wild caught nestlings is important when morphological traits are considered. This is because early- life conditions, particularly food, and maternal factors may play an important role in shaping the offspring phenotype (Dubuc-Messier *et al*., 2018; Lambert *et al*., 2021). One of the most puzzling results in this study is the fact that the city and forest birds did not differ in the F3 30 min feeding treatment, which is the standard feeding frequency used in the years the F1 and F2 birds were raised. One possible explanation is that maternal effects rather than genetics are the actual causes of the differences in F1 and F2 and they get attenuated over time. While we are unable to completely reject this possibility, we believe that the lack of difference in F3 is due to the small sample size in this group. This is supported by two additional analyses: Firstly, we have no evidence that the differences between urban and rural originating birds for the F3 generation is consistently different from the differences for the F1s and F2 birds, and if all generations are analyzed together, the results do not change (Supplements 3). Secondly, due to our egg mass results (Fig 3): differences between populations for these eggs were considerable, with city eggs being significantly lighter than those laid in forests, whereas eggs laid in aviaries by the F1 (and F2) generation did not differ between city and forest families (Fig. 3). This shows that in our study perhaps maternal factors were involved in generating city and forest differences in the size of the F1 birds. However, they should not play a role, or at least be much attenuated, in the F2 birds where size differences between city and forest were still present. Interestingly, other phenotypes such as body mass at independence (Supplements 5) were not different between city and forest birds. This lack of difference in body mass, while tarsus is larger in forest birds, is consistent with previous reports in the literature where inconsistent results have been reported for body condition (Caizergues et al., 2018, 2021). In our case, the trend is in the expected direction (Supplements 5), which could point to a genetic difference that would require a larger sample size to detect. This would makes sense given that body mass measures are more affected by the environment (e.g. time of feeding, etc) and is thus a noisier measurement.

Our experiment provides evidence that the smaller size of city birds does not allow them to better cope with a reduced availability of resources. These results are consistent with reproductive selection analyses that did not find divergent selection for body size between city and forest birds (Caizergues *et al*., 2018b). Thus, there is at present little support for the smaller size of city birds to be adaptive both in terms of reproduction and survival. Still, a smaller size could be related to adult or chick survival in ways that we did not explore. Apart from differences in the amount of food during development, urban habitats may differ from forests in terms of pollution, climate, predators, among others (Isaksson, 2018; Dominoni *et al*., 2020; Sanders *et al*., 2021). Size differences could thus impact survival of adults and/or chicks in other ways such as thermoregulation (Caizergues *et al*., 2018b) or predator avoidance (Gosler *et al*., 1995; Bitton & Graham, 2015). These hypotheses remain to be tested.

Alternatively, our experiment might not have captured the right moment of chick development for an impact to be measured. While tarsus length is a good proxy for body size (Freeman & Jackson, 1990), most of the tarsus development is already concluded by the time chicks reach 10 days old when our treatment started (Orell, 1983; Corregidor-Castro & Jones, 2021). We are confident that our treatments were effective because they did result in a difference in tarsus length between treatment groups. It is known that chicks still grow between 10 and 14-15 days old and are still susceptible to environmental effects (Corregidor-Castro & Jones, 2021) before tarsus becoming fixed. However, given our sample size, the variation and thus the power of detecting an effect would be higher if we had done the experimental manipulations at a younger age. One other methodological aspect that must be taken into account is the fact that our F3 generation consisted of birds from fewer forest populations (only Hoge Veluwe) than the F1 and F2 generations. This could have led to a more specific Hoge Veluwe versus city sites difference rather than a general difference between forests and cities. We think this aspect played a smaller role in our study compared to the sample size issue, because birds from the Hoge Veluwe population are not different in size from birds from the other forests (Supplements 2).

Finally, another plausible explanation is simply that city great tits are not adapted to the city environment (Lambert *et al*., 2021). Phenotypic differences in the urban environment are quite often assumed to be adaptive but only a handful of studies could clearly demonstrate that the distinct phenotypes lead to better survival or reproduction in their respective environments (see (Lambert *et al*., 2021) for a review). While here we clearly show genetic differences between city and forests birds they could be a result of a constant influx of small sized individuals into the population (Senar & Björklund, 2021), of founder effects or random genetic drift (Lambert *et al*., 2021). Thus, there are other non-mutually exclusive explanations for the fact that the city and forest populations differ genetically. On the one hand there is the potential that there is some degree of isolation between the populations. Studies have shown that in most locations there is gene flow between city and forest habitats, but that there is still potential for local adaptation to take place for example, via slightly reductions of gene flow along the urbanization gradient (Perrier *et al*., 2018). Another possibility is that there is non-random dispersal between city populations and surrounding rural populations (Björklund *et al*., 2010), this could lead to genetic differentiation in the absence of local adaptation (Fitzpatrick *et al*., 2017). At the present we are unable to distinguish between these different hypotheses.

In conclusion, our study shows that the smaller size of urban birds is caused by genetic differences between city and forest breeding populations rather than being only due to food contains between the city and the forest. We found no evidence for these genetic differences to be an adaptive response to lower food availability in cities.

## Author contribution

B.M.T.: conceptualization, data curation, formal analysis, funding acquisition, investigation, methodology, project administration, supervision, visualization, writing—original draft, writing—review and editing

B.T.B.: investigation, methodology, writing—review and editing; M.C.: investigation, methodology, writing—review and editing; A.W.: investigation, methodology, writing—review and editing;

K.v.O.: conceptualization, formal analysis, funding acquisition, investigation, methodology, project administration, supervision, writing—original draft, writing—review and editing; M.E.V.: conceptualization, data curation, formal analysis, funding acquisition, investigation, methodology, project administration, supervision, writing—original draft, writing—review and editing.

### Acknowledgements

We thank Barbara Rijpkema (Utrecht), Nico Tillie (Delft), Philip Kuypers (Rotterdam), Wouter Halfwerk (Amsterdam), Emily Burdfield-Steel (Amsterdam) and Henk van der Jeugd for their encouragement, support and assistance when we set-up the urban nest box sites. We thank Bart van Lith, Huib van de Haar, Ojaswi Sumbh, Maartje van Deventer, Louis Vernooij and Aurelia Strauß for their help and support during fieldwork and Anne Dijkzeul, Janina Harms, Maaike van den Born, Nina Teeuw, Ruben de Wit, Vivian van t Westende for taking good care of our birds. Finally, we thank the two referees for their comments that helped us to improve this manuscript.

## Data and code availability

Data and code underlying this article will be available in Dataverse NL upon acceptance of this manuscript (DOI https://doi.org/10.34894/XAVGM9).

