## Supplementary files for "Size in the city: morphological differences between city and forest great tits have a genetic basis"

#### 1) Seasonal changes in tarsus length

In all analyses we accounted for seasonal changes in tarsus length.

*Common garden experiment*

Although tarsus lengths vary over the season in a non-linear way ($F_{2,13.93}$ = 12.38, $p$ < 0.01, Fig. S1), birds from different origins did not differ in this pattern (origin in interaction with centered hatching date^2: $F_{1,321.75}$= 0.08, p = 0.78, origin in interaction with centered hatching date: $F_{1,244.63}$= 0.06, p = 0.80, Table S4).

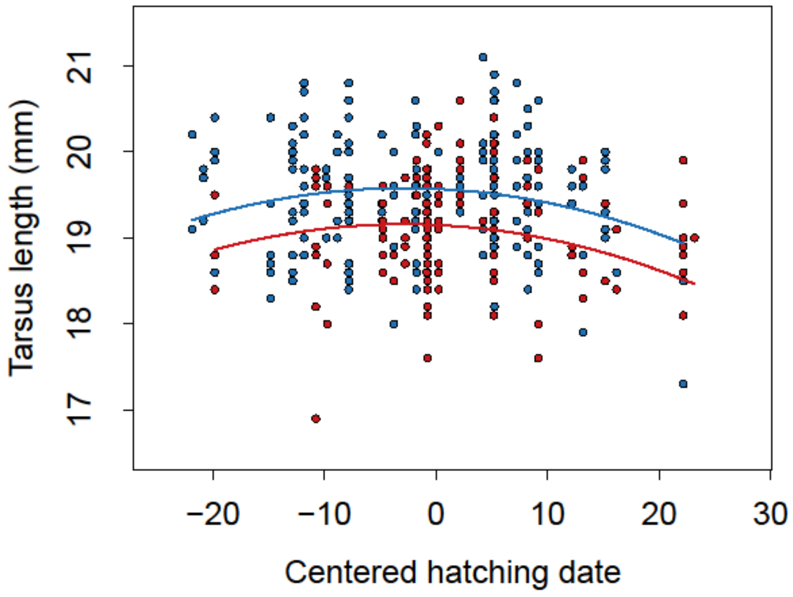


**Figure S1:** Tarsus length relative to the centered hatching date of F1 and F2 birds, points are the raw data, lines are based on the model predictions, blue = forest, red = city.

*Feeding experiment*

There was a significant interaction between origin and hatching date ($F_{1,46.25} = 10.09$, $p < 0.01$, Fig. S2, Table S5) in the presence of a hatching date squared term, indicating that the longest tarsus value was earlier in the season for city birds.

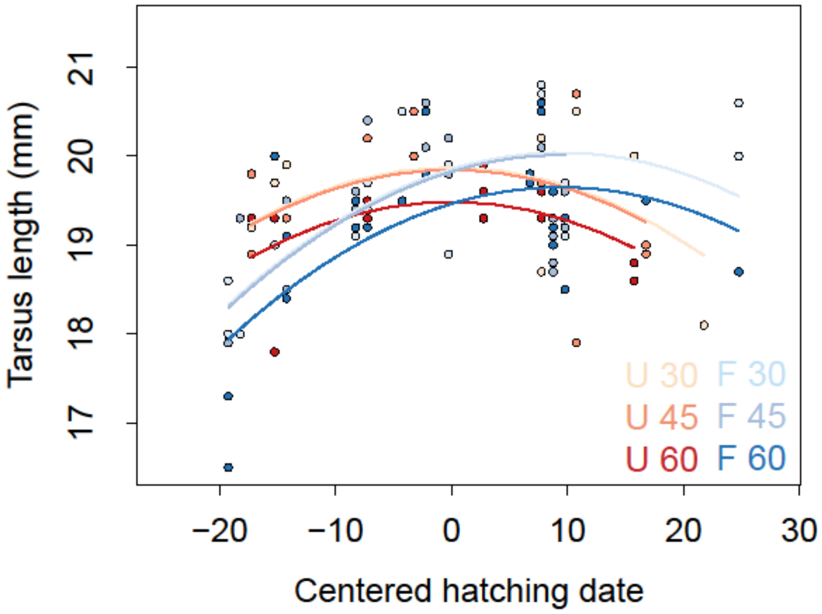


**Figure S2:** Tarsus length relative to the centered hatching date of F3 birds, points are the raw data, lines are based on the model predictions (U = City, F = Forest, 30 = food every 30 minutes, 45 = food every 45 minutes, 60 = food every 60 minutes).

Despite the limited power Fig. S2 is an interesting outcome from this experiment that we can at least speculate about. This significant interaction between origin and hatching date in the presence of a hatching squared term means that the largest tarsus values were at a different hatching date for city and forest birds, with an earlier date for city birds. This could suggest that city birds are more sensitive to low food availability earlier compared to later in the season, which would be in line with the earlier laying dates of city birds compared to forest birds in the wild (Tomotani *et al.*, 2023). This pattern could point to an adaptive early timing of breeding of city birds via the growth of the chicks. Although an interesting speculation, we must also point out that such findings were not present in the F1 and F2 years.

### 2) Comparison F1/F2 birds from Hoge Veluwe versus other forest sites

Due to logistic constraints, we only have birds from the Hoge Veluwe site as our forest population in the F3 experiment. Here we tested if there could be a difference in tarsus length of birds from the Hoge Veluwe in comparison to birds from other forest sites that could have resulted in the lack of a difference between city and forest birds in the F3 30-minute group. We fitted a model similar to all previous analyses including area (Hoge Veluwe versus other forest sites), generation (F1/F2) and sex (M/F) (centered) hatching date and (centered) HD2 as fixed effects. Family was included as random effects. We also included the interactions between origin and HD2, hatching date, sex and generation. In the model selection procedure, we dropped the interactions and main effect of area for obtaining the p-values, while retaining the terms sex, generation and hatching date/HD2 in the model as nuisance variables.

There was no difference in tarsus length between birds from the Hoge Veluwe and birds from other areas (Fig. S3), with no significant effect of area neither as a main effect ($F_{1,31.78}$= 1.04, p = 0.32) and nor in interaction with HD2 ($F_{1,201.12}$= 2.43, p = 0.12), hatching date ($F_{1,141.80}$= 2.05, p = 0.15), sex ($F_{1,196.20}$= 0.44, p = 0.51) or generation ($F_{1,32.31}$= 3.00, p = 0.09). Thus, we can assume that birds from the Hoge Veluwe are not different from birds from the other forest locations.

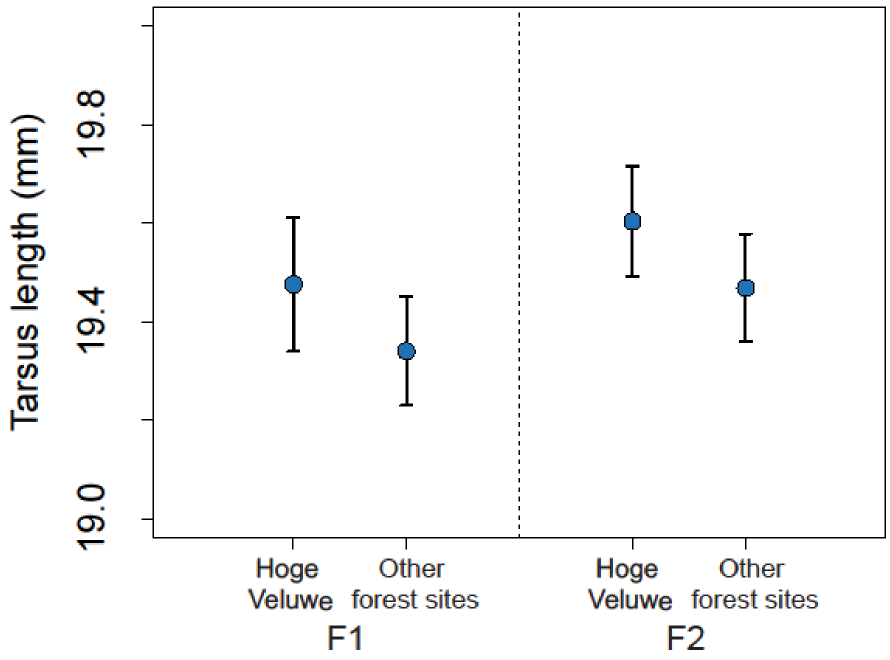


**Figure S3:** Differences in tarsus length of F1 and F2 birds from the Hoge Veluwe when compared to other forest sites. Symbols: marginal means from the models containing the additive effects of origin, sex, generation, centered hatching date^2 and centered hatching date. Bars: standard errors.

#### 3) Comparison Tarsus of F3 birds relative to F1/F2 birds

We tested if the F3 birds in the 30 min treatment (our "controls") from both origins were different from F1 and F2 birds. The same model used for comparing F1 and F2 city and forest birds was used in this test, but instead of having all 3 generations, we grouped F1 and F2 birds together to compare with F3. Thus, we included origin (city or forest), generation (F1 and F2 versus F3), sex (M/F), (centered) hatching date and (centered) HD2 as fixed effects and family nested within population as random effect. We also included the interactions between origin and HD2, hatching date, sex and generation. In the model selection procedure, we dropped the interactions and main effect of origin and generation for obtaining the p-values, while retaining the terms sex and hatching date/HD2 in the model as nuisance variables.

The inclusion of F3 "control" birds did not change the results obtained when comparing city and forest birds using only F1 and F2 birds. There was again a non-linear change in tarsus length over the season ($F_{1,348.24}$ = 12.32, $p < 0.01$, Fig. S4), with birds from different origins not differing in this pattern (origin in interaction with centered hatching date^2: $F_{1,346.86}$ = 0.003, p = 0.96, origin in interaction with centered hatching date: $F_{1,270.27}$= 0.04, p = 0.83). Birds originating from cities once again had a smaller tarsus ($F_{1,3.87}$= 10.06, p = 0.04, city estimate = 18.50 ±0.12mm, forest estimate = 18.85 ±0.11mm). Generation (F1 and F2 versus F3) did not have an effect on tarsus length neither as main effect ($F_{1,76.12}$= 1.70, p = 0.20) and nor in interaction with origin ($F_{1,122.13}$= 0.83, p = 0.37).

We can't completely exclude the possibility that the differences we found between city and forest birds in F1 and F2 are pure maternal effects that weaken over time. However, the results of this additional analysis suggest that it is more likely that the reduced sample size of F3 (as can also be seen in Fig. S4) played a stronger role than a possible biological difference between F3 and F1/F2.

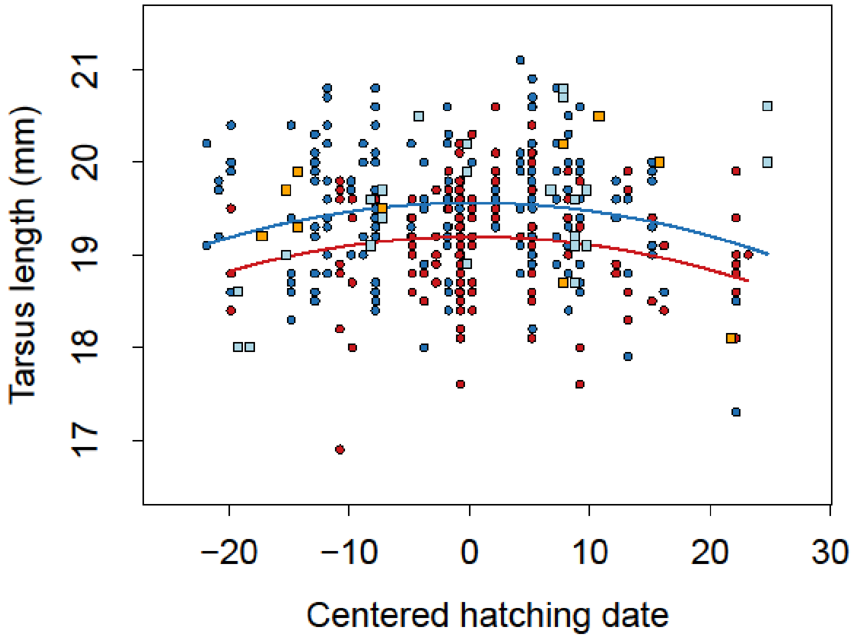


**Figure S4:** Tarsus length relative to the centered hatching date of F1, F2 and F3 birds, points are the raw data, lines are based on the model predictions, circles = F1 and F2 birds, blue = forest, red = city; squares = F3 birds in the 30-min treatment, light blue = forest, orange = city.

### 4) Hatching date and body mass common garden experiment

All chicks were weighed when they arrived at the institute with 10 days old, then a second time at independence at around 32-40 days old when they were feeding on their own

In three separate analyses, we compared if a) the hatching dates, b) the body masses when 10 days old (the time when they were taken from the field to the aviaries) and c) the body masses at independence differed between birds with a genetic city or forest origin using birds from the F1 and F2 generations. We included origin (city or forest), generation (F1/F2) and sex (M/F) as fixed effects. In the case of the body mass measurements, we also included hatching date and hatching date squared as fixed effects in order to correct for differences in the foster brood condition, which were centered around the mean hatching date per year. To take into account potential genetic differences of family or population of origin, we included family nested within population as random effects. We also included the interactions between origin and sex and generation, in the case of fledge age and body masses, we also included the interaction between origin, HD2 and hatch date. In the model selection procedure, we dropped the interactions and main effect of origin for obtaining the p-values, while retaining the terms sex, generation and hatching date/HD2 in the model as nuisance variables.

There was a significant interaction between origin and generation for hatching date ($F_{1,57.75}$ = 23.67, $p$ <0.01, Table S4), with birds from different origins and generations differing in their hatching dates (Fig. S5a, Table S4). There was no difference in weight between birds from city and forest families, neither in their weight when 10 days old ($F_{1,6.21}$ = 0.06, $p$ = 0.82, Fig. S5b, Table S4), nor in their weight at independence ($F_{1,6.11}$ = 1.85, $p$ = 0.22, Table S4) although the latter showed a trend in the expected direction (Fig. S5c, Table S4).

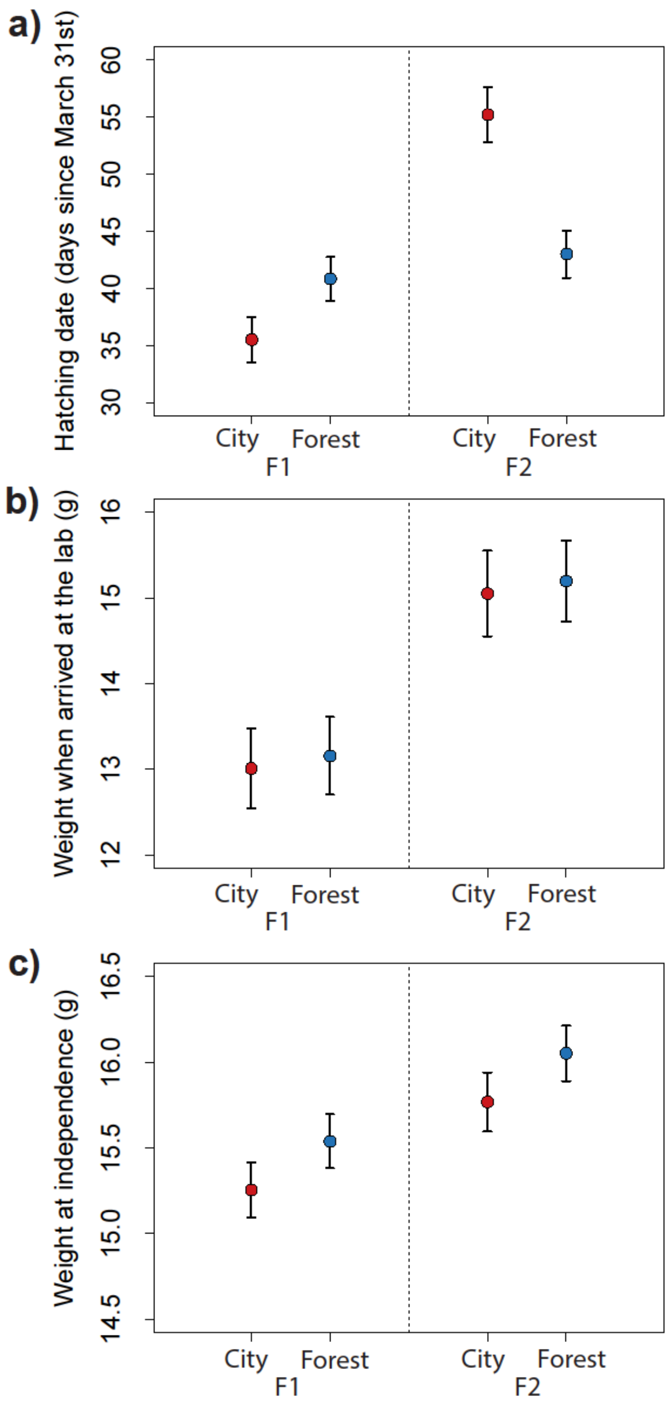


**Figure S5: a)** Differences in hatching date of F1 and F2 birds. **b)** Differences in body mass at arrival to the lab of F1 and F2 birds. **c)** Differences in body mass at independence of F1 and F2 birds. Closed symbols: marginal means from the models containing the additive effects of origin, sex, generation, centered hatching date^2 and centered hatching date. Bars: standard errors; open symbols: raw measurements.

### 5) Body mass feeding experiment

Similarly to the F1 and F2 chicks, birds of the third generation (F3) were weighed twice, once when they arrived at the institute with 10 days old and a second time at independence at around 32-40 days old. Specifically in this feeding experiment, chicks were also weighed an additional time at fledging which marked the end of the experimental feeding regime.

We tested if birds from distinct origins and in different treatments had a different fledge age or body mass at fledging and once independent. We fitted treatment (30 min, 45 min or 60 min), origin (city or forest) and sex (M/F) as fixed effects and also included the hatching date and HD2 as fixed effects like in the previous analysis. Once more, to account for potential genetic differences of family or population of origin, we included family nested within population as random effects. Apart from the interaction between origin and treatment we also included the interactions between origin and hatch date, origin and sex, treatment and hatch date and treatment and sex. In the model selection procedure, we dropped the interactions and main effect of origin and treatment for obtaining the p-values, while retaining the terms sex, generation and hatching date/HD2 in the model as nuisance variables.

Birds originating from city or forest families did not respond differently to the treatments (no significant interaction between origin and treatment) and thus city and forest birds did not differ in their fledge weight ($F_{2,75.42}$ = 0.87, $p$ =0.42, Table S5) or weight at independence ($F_{2,79.78}$ = 0.35, $p$ =0.26, Table S5) (Fig. S6b,c, Table S5). There was also no overall treatment effect on the fledging weight ($F_{2,80.89}$ = 2.22, p = 0.12, Fig. S6a, Table S5) or weight at independence ($F_{2,83.76}$ = 2.17, p = 0.12, Fig. S6b, Table S5), thus birds in different treatments did not differ in their body mass.

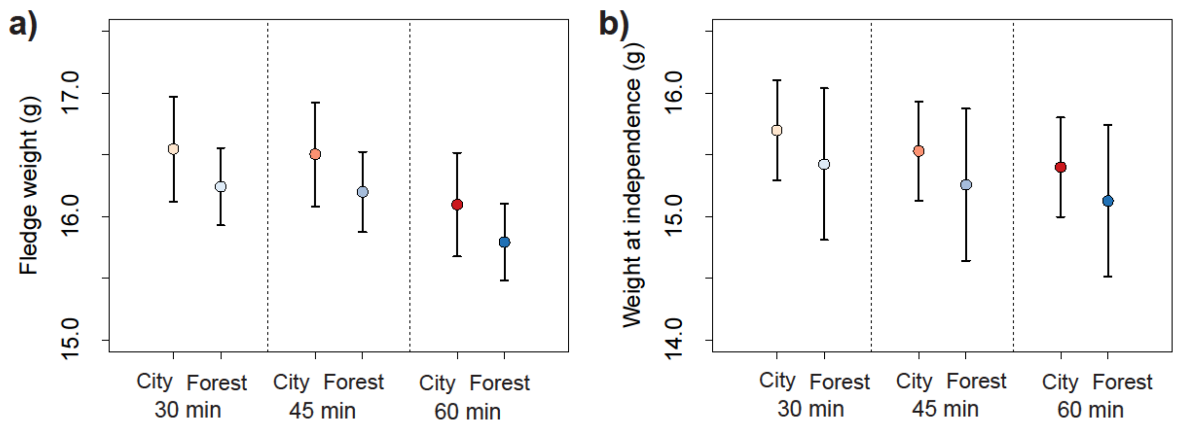


**Figure S6: a)** Differences in body mass at fledging of F3 birds. **b)** Differences in body mass at independence of F3 birds. Closed symbols: marginal means from the models containing the additive effects of treatment, origin, sex, generation, centered hatching date^2 and centered hatching date. Bars: standard errors; open symbols: raw measurements.

### 6) Supplementary tables

**Table S1: Detailed sample sizes of each step of the study.**

| Type | Location | Sex | Wild breeding | Common garden (F1) | Common garden (F2) | Feeding experiment (F3) |
|---|---|---|---|---|---|---|
| Forest | Heikamp (near Hoge Veluwe) | F | 26 | | | |
| Forest | Heikamp (near Hoge Veluwe) | M | 16 | | | |
| Forest | Hoge Veluwe National Park | F | | 12 | 31 | 36 |
| Forest | Hoge Veluwe National Park | M | | 13 | 31 | 30 |
| Forest | Bennekom | F | | 3 | | |
| Forest | Bennekom | M | | 7 | | |
| Forest | Liesbos | F | | 1 | | |
| Forest | Liesbos | M | | 2 | | |
| Forest | Oosterhout | F | | 7 | 11 | |
| Forest | Oosterhout | M | | 5 | 15 | |
| Forest | Vlieland | F | | 9 | 27 | |
| Forest | Vlieland | M | | 12 | 27 | |
| Forest | Total | | 42 | 71 | 142 | 66 |
| City | Amsterdam | F | | 9 | 12 | 4 |
| City | Amsterdam | m | | 9 | 14 | 7 |
| City | Delft | f | | 7 | 9 | 6 |
| City | Delft | m | | 6 | 6 | 6 |
| City | Rotterdam | f | | 4 | 5 | 7 |
| City | Rotterdam | m | | 6 | 3 | 4 |
| City | Utrecht | f | 22 | 14 | 10 | |
| City | Utrecht | m | 31 | 6 | 1 | |
| City | Total | | 53 | 61 | 60 | 34 |

**Table S2: Comparison of the tarsus length of wild caught city and forest birds.** Effects of site (city or forest), sex (male or female) and the interaction between the two. Main term capture year is included in the model but not tested. Statistics are given at the point of exclusion of the term from the model, significant effects are given in bold.

| **Tarsus length** | **Estimate** | **s.e.** | **Ndf** | **Ddf** | ***F*-value** | ***p*-value** |
|---|---|---|---|---|---|---|
| site:sex | | | 1.00 | 57.81 | 0.43 | 0.51 |
| Site | | | 1.00 | 48.18 | 5.76 | **0.02** |
| Sex | | | 1.00 | 41.17 | 20.27 | **< 0.01** |
| site (city) | 18.39 | 0.45 | | | | |
| site (forest) | 18.72 | 0.45 | | | | |
| sex (male) | 0.61 | 0.12 | | | | |
| Year (2019) | 0.50 | 0.47 | | | | |
| Year (2020) | 0.72 | 0.45 | | | | |
| Year (2021) | 0.88 | 0.47 | | | | |
| Year (2023) | 1.13 | 0.51 | | | | |

**Table S3: Comparison of the egg masses of city and forest birds laid by females from the P (wild), F1 and F2 (aviaries) generations.** Effects of site (city or forest), mother generation (P1, F1, F2) and the interaction between the two. Main term of laying date is included in model but not tested. Statistics are given at the point of exclusion of the term from the model, significant effects are given in bold.

| **Egg mass** | **Estimate** | **s.e.** | **ndf** | **ddf** | **F-value** | **p-value** |
|---|---|---|---|---|---|---|
| mother origin:mother generation | | | 2.00 | 80.77 | 2.94 | **0.06** |
| mother origin | | | 1.00 | 3.91 | 3.98 | 0.12 |
| mother generation | | | 2.00 | 102.97 | 11.66 | **< 0.01** |
| mother origin (city) | 166.11 | 2.81 | | | | |
| mother origin (forest) | 170.08 | 2.15 | | | | |
| mother generation (F2) | -0.04 | 4.76 | | | | |
| mother generation (P) | -18.07 | 3.97 | | | | |
| laying date | 0.17 | 0.01 | | | | |
| mother origin (forest) : mother generation (F2) | -6.17 | 5.77 | | | | |
| mother origin (forest) : mother generation (P) | 8.99 | 5.10 | | | | |
| **Post hoc egg mass** | **estimate** | **s.e.** | | **df** | **t.ratio** | **p.value** |
| P (city x forest) | -12.96 | 4.13 | | 25.20 | -3.14 | **< 0.01** |
| F1 (city x forest) | -3.97 | 3.78 | | 17.40 | -1.05 | 0.31 |
| F2 (city x forest) | 2.20 | 5.60 | | 13.10 | 0.39 | 0.70 |

**Table S4: Comparison between birds originating from city and forest families (F1 and F2 generations).** Effects of origin and interaction of origin and generation (a,b) and origin and hatching date^2, hatching date, sex and generation (c,d,e,f) on the a) hatching date ^2, b) hatching date, c) weight when 10 days old, d) weight at independence and e) tarsus length. Main terms of sex, generation, hatching date and hatching date^2 are included in the models but not tested. Statistics are given at the point of exclusion of the term from the model, significant effects are given in bold.

| **a) Hatching date^2** | **Estimate** | **s.e.** | **ndf** | **ddf** | ***F*-value** | ***p*-value** |
|---|---|---|---|---|---|---|
| origin:generation | | | 1.00 | 57.76 | 23.66 | **< 0.01** |
| origin (city) | 1295.84 | 175.82 | | | | |
| origin (forest) | 1718.03 | 169.48 | | | | |
| generation (F2) | 1869.49 | 249.32 | | | | |
| sex (male) | -33.60 | 64.31 | | | | |
| origin (forest):generation (F2) | -1582.19 | 322.23 | | | | |
| **b) Hatching date** | **Estimate** | **s.e.** | **ndf** | **ddf** | ***F*-value** | ***p*-value** |
| origin:generation | | | 1.00 | 57.75 | 23.67 | **< 0.01** |
| origin (city) | 35.63 | 2.00 | | | | |
| origin (forest) | 40.95 | 1.92 | | | | |
| generation (F2) | 19.71 | 2.76 | | | | |
| sex (male) | -0.25 | 0.69 | | | | |
| origin (forest):generation (F2) | -17.53 | 3.57 | | | | |
| **c) Weight when 10 days old** | **Estimate** | **s.e.** | **ndf** | **ddf** | ***F*-value** | ***p*-value** |
| origin:hatch date^2 | | | 1.00 | 319.25 | 1.22 | 0.27 |
| origin:hatch date | | | 1.00 | 264.31 | 1.99 | 0.16 |
| origin:generation | | | 1.00 | 65.79 | 1.03 | 0.31 |
| origin:sex | | | 1.00 | 303.97 | 0.09 | 0.77 |
| Origin | | | 1.00 | 6.21 | 0.06 | 0.82 |
| centred hatch date squared | -0.01 | 0.00 | | | | |
| centred hatch date | -0.08 | 0.01 | | | | |
| origin (city) | 13.36 | 0.47 | | | | |
| origin (forest) | 13.51 | 0.46 | | | | |
| generation (F2) | 2.04 | 0.38 | | | | |
| sex (male) | 0.36 | 0.19 | | | | |
| **d) Weight at independence** | **Estimate** | **s.e.** | **ndf** | **ddf** | ***F*-value** | ***p*-value** |
| origin:hatch date^2 | | | 1.00 | 318.90 | 0.68 | 0.41 |
| origin:hatch date | | | 1.00 | 239.25 | 0.15 | 0.70 |
| origin:generation | | | 1.00 | 62.95 | 1.30 | 0.26 |
| origin:sex | | | 1.00 | 310.60 | 0.19 | 0.66 |
| Origin | | | 1.00 | 6.11 | 1.85 | 0.22 |
| centred hatch date squared | 0.00 | 0.00 | | | | |
| centred hatch date | -0.01 | 0.01 | | | | |
| origin (city) | 14.99 | 0.16 | | | | |
| origin (forest) | 15.27 | 0.16 | | | | |
| generation (F2) | 0.51 | 0.14 | | | | |
| sex (male) | 0.87 | 0.08 | | | | |

| e) Tarsus length | Estimate | s.e. | ndf | ddf | *F*-value | *p*-value |
|---|---|---|---|---|---|---|
| origin:hatch date^2 | | | 1.00 | 321.75 | 0.08 | 0.78 |
| origin:hatch date | | | 1.00 | 244.63 | 0.06 | 0.80 |
| origin:generation | | | 1.00 | 64.05 | 2.18 | 0.14 |
| origin:sex | | | 1.00 | 311.75 | 0.66 | 0.42 |
| Origin | | | 1.00 | 5.67 | 9.02 | **0.03** |
| centred hatch date squared | 0.00 | 0.00 | | | | |
| centred hatch date | -0.01 | 0.00 | | | | |
| origin (city) | 18.86 | 0.10 | | | | |
| origin (forest) | 19.20 | 0.09 | | | | |
| generation (F2) | 0.27 | 0.11 | | | | |
| sex (male) | 0.45 | 0.06 | | | | |

**Table S5: Results of the experiment where birds from city and forest families (F3) were fed at different frequencies.** Effects of origin, treatment and the interaction between them, hatching date and sex on a) fledge weight, b) weight upon independence and c) tarsus length. The main effect of treatment and origin was also tested in a separate analysis when the interaction was significant.

| **a) Fledge weight** | **Estimate** | **s.e.** | **ndf** | **ddf** | ***F*-value** | ***p*-value** |
|---|---|---|---|---|---|---|
| origin:treatment | | | 2.00 | 75.42 | 0.87 | 0.42 |
| origin:hatch date | | | 1.00 | 42.91 | 1.49 | 0.23 |
| treatment:hatch date | | | 2.00 | 72.79 | 0.32 | 0.73 |
| origin:sex | | | 1.00 | 78.61 | 0.48 | 0.49 |
| treatment:sex | | | 2.00 | 79.08 | 1.79 | 0.17 |
| Origin | | | 1.00 | 0.26 | 0.39 | 0.78 |
| Treatment | | | 2.00 | 80.99 | 2.22 | 0.12 |
| centred hatch date squared | 0.00 | 0.00 | | | | |
| centred hatch date | 0.03 | 0.01 | | | | |
| treatment (30 min) | 16.53 | 0.45 | | | | |
| treatment (45 min) | 16.49 | 0.44 | | | | |
| treatment (60 min) | 16.08 | 0.43 | | | | |
| origin (forest) | -0.30 | 0.46 | | | | |
| sex (male) | 0.92 | 0.20 | | | | |
| **b) Weight at independence** | **Estimate** | **s.e.** | **ndf** | **ddf** | ***F*-value** | ***p*-value** |
| origin:treatment | | | 2.00 | 79.78 | 0.35 | 0.26 |
| origin:hatch date | | | 1.00 | 71.02 | 0.72 | 0.40 |
| treatment:hatch date | | | 2.00 | 75.43 | 0.72 | 0.49 |
| origin:sex | | | 1.00 | 84.17 | 3.14 | 0.08 |
| treatment:sex | | | 2.00 | 81.59 | 2.30 | 0.11 |
| Origin | | | 1.00 | 1.43 | 0.14 | 0.75 |
| Treatment | | | 2.00 | 83.76 | 2.17 | 0.12 |
| centred hatch date squared | 0.00 | 0.00 | | | | |
| centred hatch date | 0.02 | 0.01 | | | | |
| treatment (30 min) | 15.49 | 0.42 | | | | |
| treatment (45 min) | 15.33 | 0.41 | | | | |
| treatment (60 min) | 15.20 | 0.41 | | | | |
| origin (forest) | -0.27 | 0.72 | | | | |
| sex (male) | 0.87 | 0.12 | | | | |
| **c) Tarsus length** | **Estimate** | **s.e.** | **ndf** | **ddf** | ***F*-value** | ***p*-value** |
| origin:treatment | | | 2.00 | 74.12 | 0.07 | 0.93 |
| origin:hatch date | | | 1.00 | 46.25 | 10.09 | **< 0.01** |
| treatment:hatch date | | | 3.00 | 77.72 | 1.56 | 0.21 |
| origin:sex | | | 1.00 | 79.86 | 0.50 | 0.48 |
| treatment:sex | | | 2.00 | 75.32 | 0.14 | 0.87 |
| Treatment | | | 2.00 | 80.01 | 3.91 | **0.02** |
| centred hatch date squared | 0.00 | 0.00 | | | | |
| centred hatch date | -0.01 | 0.01 | | | | |
| treatment (30 min) | 19.54 | 0.25 | | | | |
| treatment (45 min) | 19.54 | 0.25 | | | | |

| | | |
|---|---|---|
| treatment (60 min) | 19.25 | 0.24 |
| origin (forest) | 0.06 | 0.27 |
| sex (male) | 0.51 | 0.10 |
| centred hatch date:origin (forest) | 0.05 | 0.01 |
